# Seeing the future: connectome strength and network efficiency in visual network predict individual ability of episodic future thinking

**DOI:** 10.1101/2022.10.06.511122

**Authors:** Bowen Hu, Rong Zhang, Tingyong Feng

## Abstract

Episodic future thinking (EFT) refers to the critical ability that people construct vivid mental imagery about future events and pre-experience it, which helps with individual and group decision-making. Although EFT is generally believed to have a visual nature by theorists, little neuroscience evidence had been provided to verify this assumption. Here, by employing the approach of connectome-based predictive modeling (CPM) and graph-theoretical analysis, we analyzed resting-state functional brain image from 191 participants to predict their variability of EFT ability (leave-one-out cross-validation), and validated the results by applying different parcellation schemas and feature selection thresholds. At the connectome strength level, CPM-based analysis revealed that EFT ability could be predicted by the connectome strength of visual network. Further at the network level, graph-theoretical analysis showed that EFT ability could be predicted by the network efficiency of visual network. Moreover, these findings were replicated using different parcellation schemas and feature selection thresholds. These results robustly and collectively supported the visual network being the neural substrates underlying EFT ability from a comprehensive perspective of resting-state functional connectivity strength and the neural network. This study provides indications on how the function of visual network supports EFT ability, and helps to understand the EFT ability from a neural basis perspective.

## 1 Introduction

People often think ahead about events that are going to happen in the future and construct vivid scenes of them, to pre-experience and evaluate what might be happening, such as an upcoming vacation or a low-carbon future. This ability is called episodic future thinking (EFT), which is also known as foresight (Peters & Büchel, 2010). EFT consists of two processes, constructing vivid scenes of future events and subjectively experiencing the constructed scenes (D’Argembeau & Van der Linden, 2012). EFT has a beneficial influence on intertemporal decision-making such as saving money for the future or health-related behavior (Peters & Büchel, 2010; Rösch et al., 2021), and on psychological well-being by simulating more episodic detail for constructive behaviors (Jing et al., 2016). Yet such an ability varies across people, for example, some individuals can’t construct future scenes at all (for example aphantasia), but some others can construct future scenes that are photo-like (Zeman et al., 2020). Thus, some people with good EFT ability may envision and make better choices for the future, while the others lose sight of the future and cannot make the right choice. This diversity may also emerge on the group or social level such as ill-considered policies issued by the nearsighted government, which may undermine well-being for mankind. Considering the beneficial impact of EFT on improving individual and collective well-being, it is significant to explore the neural substrates underlying the EFT ability, which may help us understand the neural basis of EFT ability and predict it from the neuroscience perspective.

The default mode network (Raichle, 2015) is considered to play a role in EFT, especially the scene construction process(Østby et al., 2012; Palombo et al., 2018; Pearson, 2019; Stawarczyk & D’Argembeau, 2015). Activation of DMN regions including lateral parietal cortex, ventromedial prefrontal cortex (vmPFC), hippocampus, and posterior cingulate cortex (PCC) has been observed when imagining future events(Gaesser et al., 2013; Madore et al., 2016; Palombo et al., 2018; Szpunar et al., 2007; van Mulukom et al., 2013). More importantly, functional connectivity in DMN is found to be related to the EFT ability in adolesœnts(Østby et al., 2012). In theory, the relationship between EFT and DMN can be understood through the concept of mental time travel, which referred to individuals’ re-experiencing the past or pre-experiencing the future(Bar, 2009; Boyer, 2008; Østby et al., 2012). The ability of mental time travel is largely dependent on the function of DMN for self-projecting oneself into the situation, subsequently constructing the past or future episodes(Buckner & Carroll, 2007; Hassabis & Maguire, 2007; Østby et al., 2012). Thus, DMN may serve as a potential neural substrate for the EFT ability, especially the ability to subjectively experience the constructed scenes.

Apart from DMN, there is also mounting evidence suggesting that the variation of EFT ability may also come from the difference in the visual function and the related visual network. Firstly, in a behavioral experiment, when subjects were instructed to focus their sight on a moving dot(guided eye movement) during EFT, it would significantly impair the episodic details generated compared with the control condition(de Vito et al., 2015), and behavioral researchers also found that an individual’s visual spatial imagery ability can predict its EFT ability (Aydin, 2018). Moreover, neuroscientist also suggest that visual network plays an important role in EFT. For example, Conti and Irish(2021) proposed that the functional coupling between the visual cortex and DMN will flexibly extract sensory-perceptual representations from episodic memory to support future thinking, while Pearson (2019) also proposed a similar idea that imagery content retrieves from DMN will be formed in the visual cortex. These ideas are supported by neuroscience research evidence, for example, neural activity in the visual cortex is important for information encoding during mental imagery (Dijkstra et al., 2017; Naselaris et al., 2015; Quoidbach et al., 2008), and the functional connectivity between the hippocampus (which is a part of DMN) and the visual cortex increased significantly during EFT (Bellana et al., 2016). Taken together, the evidence above suggests that the visual function and the neural activity in the visual cortex play an essential role in EFT, thus the neural substrates underlying the variation of EFT ability may lie in the difference of visual network function. Yet there is no known research to help understand or predict this variation from the neuroscience perspective, such as resting-state functional connectivity (rsFC) and visual neural network.

RsFC is the temporal correlation between neural activities of brain regions, reflecting stable information integrations across brain regions (Faria et al., 2012; van den Heuvel & Hulshoff Pol, 2010), thus rsFC is an ideal way to capture the neural substrates underlying complex cognitive process, personality and ability (Beaty et al., 2018; Finn et al., 2015; Nostro et al., 2018). Connectome-based predictive modeling (CPM) utilizes all the functional connection strength within the whole-brain network or subnetwork to predict certain ability, revealing the neural substrates underlying psychological traits from both whole-brain network or a functional subnetwork perspective (Finn et al., 2015; Shen et al., 2017). CPM has been widely and successfully used to predict fluid intelligence, creativity ability, internet gaming disorder and mnemonic discrimination (Beaty et al., 2018; Shen et al., 2017; Song et al., 2021; Wahlheim et al., 2021), therefore being an appropriate method for exploring the association between intrinsic functional connectivity and EFT ability.

With the brain being a complex system, only connection strength cannot capture the whole picture of brain functions and characteristics, because brain functions depend not only on the simple summation of the functional connectivity (Finn et al., 2015; Shen et al., 2017), but also largely on the integrated network organization of the brain (Bullmore & Sporns, 2012a; Haimovici et al., 2013; Pessoa, 2014). Thus, the intrinsic topologic efficiency of brain network (like network efficiency) is fundamental for brain functioning and often regarded as neural markers for psychological traits (Bullmore & Sporns, 2012a; Chen et al., 2018; Pineda-Pardo et al., 2016).

The present study aimed to investigate the neural substrates responsible for EFT ability and to predict variation of EFT ability, especially focusing on the ability to generate episodic details. To achieve these aims, we would focus on the analysis of the intrinsic functional connectome strength and the topological property (network efficiency) of brain network, using CPM analysis and graph-theoretical analysis to make prediction (leave-one-out cross-validation) of EFT ability. And based on the previous analysis, we hypothesized that visual network and DMN are the neural substrates that accounted for the variation of EFT ability.

## 2 Materials and Methods

### 2.1 Participants and procedure

The current study recruited 196 students (46 males; M = 19.99 years old) from Southwest University, China. Five participants were excluded from further analysis due to excessive head motion (Jenkinson et al., 2002), leaving 191 participants for further analyses. All participants were right-handed and had no history of psychiatric disorder or drug abuse. This study was proved by the Institutional Review Board of Southwest University. All participants gave informed consent and reported their demographic information before they completed EFT measure task and MRI scan. After the experiment, participants were paid for their participation.

### 2.2 EFT measure task

In this study, we applied a questionnaire from the paradigm developed by D’Argembeau & Linden (2012) to measure EFT ability. EFT consisted of two dimensions, Sensory-perceptual Qualities (EFT-SPQ) and Autonoetic Consciousness (EFT-AC). EFT-SPQ measures how vivid the constructed future scenes are, and EFT-AC measures how deeply one got involved in the future thinking. Participants were instructed to imagine a series of events based on cue words, which were all daily events that may happen to participants in real life (i.e., family, study, friend, party, and trip), and participants were instructed to imagine events happening at least one year later. All the cue words were presented individually to each participant and the order of the cues was counterbalanced across participants. To ensure that participants fully imagined future events, we asked them to think of each event for longer than one minute and try to keep the imagination in mind (D’Argembeau & Van der Linden, 2012). Right after completing each event, participants filled a 7-point Likert-type questionnaire assessing the sensory-perceptual qualities (D’Argembeau & Van der Linden, 2012), including overall vividness (1 = vague, 7 = extremely vivid), amount of visual details (1 = not at all, 7 = a lot), amount of other sensory details (1 = not at all, 7 = a lot), clarity of imagined location (1 = not at all clear, 7 = extremely clear), and clarity of persons and objects (1 = very vague, 7 = very clear). To ensure the validity of this measure, each item was explained carefully to the participants. We calculated the summed scores of all five items as the score of EFT-SPQ. The Cronbach alpha coefficient of EFT-SPQ of current data was 0.91, which indicates excellent reliability. The other dimension of EFT-AC was also assessed together with EFT-SPQ, but this dimension was not the focus of this study, thus received no further analysis.

### 2.3 fMRI data acquisition and preprocessing

The resting-state fMRI images were acquired on a TRIO 3.OT scanner (Siemens Medical, Erlangen, Germany). High-resolution T1 structural images were acquired using magnetization prepared rapid acquisition gradient-echo (MPRAGE) sequence (TR/ TE = 2530ms/ 3.39ms; slices = 128; flip angle = 7°; matrix size = 256 × 256; voxel size = 1×1 × 1.33 mm3). Resting-state functional images were acquired using echo-planar imaging sequence (TR/TE = 2,000ms/ 30ms; slices = 33; flip angle = 90°; FOV = 200× 200; resolution matrix = 64× 64× 33; voxel size =3.1 × 3.1 × 3.6 mm^3^).

The resting-state functional images were preprocessed using Data Processing Assistant for resting-state fMRI (Yan & Zang, 2010). The first 10 volumes were discarded because of the 200; effect of magnetization disequilibrium and the adaptation of subjects to the circumstance, then slice-timing correction and realignment were applied to the remaining volumes. Next, the functional images were spatially normalized to the MNI space (3 × 3 × 3 mm^3^ voxel sizes) and smoothed with a 4mm full-width-at-half-maximum (FWHM) Gaussian kernel. Finally, the nuisance signals (including WM, CFS, global signal, Friston 24 parameters) were regressed out, temporal filtering (0.01–0.08 HZ) and linear detrending were performed.

### 2.4 Functional network construction

To generate functional connectivity matrices of DMN and visual network, first, we defined the network nodes by using Power 264-node functional brain atlas (Power et al., 2011). Then the DMN and visual network were constructed followed by the grouping schemas proposed by Power et al. (2011). Pearson’s correlation coefficients were calculated on mean time courses (i.e., the average time course of all the voxels within the node) of each paired nodes, which finally generated symmetric adjacent matrices for each participant. The elements of the matrix served as edge strengths in the network analysis.

### 2.5 Connectome-based predictive modeling

CPM analysis was conducted by MATLAB (https://www.mathworks.com) scripts modified from the original CPM scripts provided by Shen et al. (2017). In each iteration of leave-one-out cross-validation (LOOCV) in model training and testing stage, to avoid the arbitrary choice of feature selection threshold, we implemented three different thresholds (significance *p* = 0.1, 0.05, and 0.01, the thresholds are same as in Finn et al. (2015)) to converge on a robust result. Specifically, connectomes that were poorly correlated with the behavior measure (i.e., the significance of the correlation coefficients were above threshold) in the training set would be set to zeros. Besides, those connectomes that were significantly positively or negatively correlated with the behavior measure will be summed into one value for each subject respectively, namely positive network strength and negative network strength, which was used as a predictor to predict behavioral measure in a linear model. Then in the model testing stage, the connectivity mask and parameter of predictive model acquired in the training stage would be applied to predict the behavioral score of the left-out subject, based on the summed connectome strength. After all the iterations of cross-validation were done, the correlation coefficients of the predictive score and the real score of the subjects would be calculated to assess model performance.

The process above was applied to whole-brain network and 14 subnetworks defined by Power and his colleagues (2011). To further verify the reliability of results, we also applied two other brain functional atlas with varying scales (each including 400 and 1024 regions) to rule out the potential effects of parcellation schemas (Schaefer et al., 2018; Zalesky et al., 2010). The processing steps were same as above-mentioned. Finally, we performed a permutation test to assess the significance of our predictive model. To be more specific, by randomly shuffling the corresponding behavioral score of the subjects for 1000 iterations, in each iteration we reran all the CPM analysis mentioned above and assess the null-model performance. The *p* value is defined as the probability that the performance of the real model exceeds the null-models, which indicates the significance of the real model’s prediction performance.

### 2.6 Network topology analysis of the network

To quantify the topological efficiency of the network, we calculated global efficiency (*E_glob_*) for the whole-brain network and subnetwork (Latora & Marchiori, 2001). First, we removed the spurious correlation in the acquired adjacent matrix by applying a threshold of 0.2, and then the remaining edges (adjacencies) were used to generate weighted functional networks. Finally, for each network, we calculated *E_glob_* which quantified the efficiency of the parallel information transfer among all possible pairs of nodes in the network *G*, defined as follows:

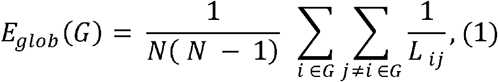

where *L_ij_* is the shortest path length between node *i* and node *j* in the network *G*, and *N* denoted the number of nodes in a certain network (e.g., N = 33 in the visual network in Power atlas). Network topology analysis was conducted in Graph Theoretical Network Analysis (GRETNA) Toolbox (Wang et al., 2015). We ran the same linear prediction model with leave-one-out cross-validation as we did in 2.4, in which the predictor became *E_glob_* and the outcome value was still EFT-SPQ. Considering the potential effects of the correlation threshold on the analysis, we additionally applied two other thresholds (0.1 and 0.3) to validate our result.

## 3 Results

### 3.1 Behavioral results

The results showed that EFT-SPQ scores were normally distributed (Kolmogorov-Smirnov statistical magnitude = 0.062, *p*= 0.073, see Figure 1), the average score of EFT-SPQ was 117.70, and the *SD* was 18.44.

**FIGURE 1.**
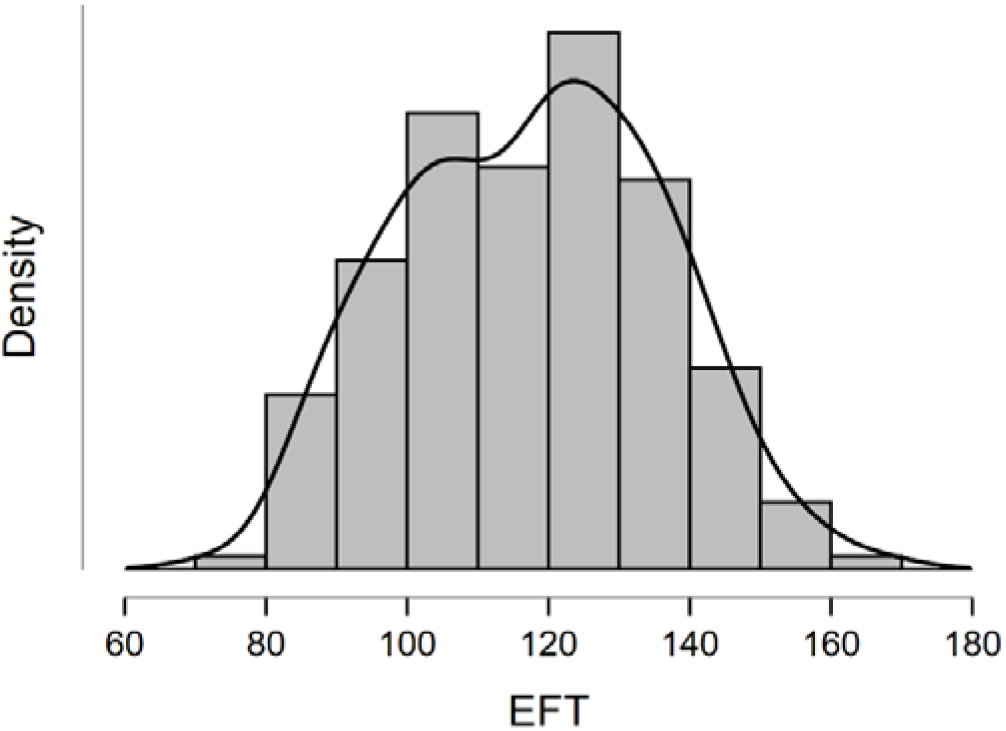
Distribution of EFT.

### 3.2 Connectome-based predictive modeling

We performed CPM on the proposed DMN and visual subnetworks, and results showed that positive prediction model of EFT generated from visual network connectivity was significant (using 0.1, 0.05, and 0.01 as feature selection thresholds, *r*(191) = 0.24, 0.25 and 0.24, p < 0.001; for results of 0.05 threshold, see Figure 2A). The connections that contributed to the significant prediction model are displayed in Figure 2B. These findings were consistent across all feature selection thresholds and parcellation schemas were used (when applying Schaefer-400 atlas, using 0.1, 0.05, and 0.01 as feature selection thresholds, *r*(191) = 0.18, 0.18 and 0.20, p < 0.05; when applying Zalesky-1024 atlas, using 0.1, 0.05, and 0.01 as feature selection thresholds, *r*(191) = 0.19, 0.20 and 0.19, p < 0.01; see Table 1). Collectively these results robustly indicated that intrinsic visual network connectivity can predict EFT-SPQ, thus could be considered the neural substrate of EFT ability, which supported our hypothesis that visual network is the neural substrate underlying EFT ability.

**FIGURE 2.**
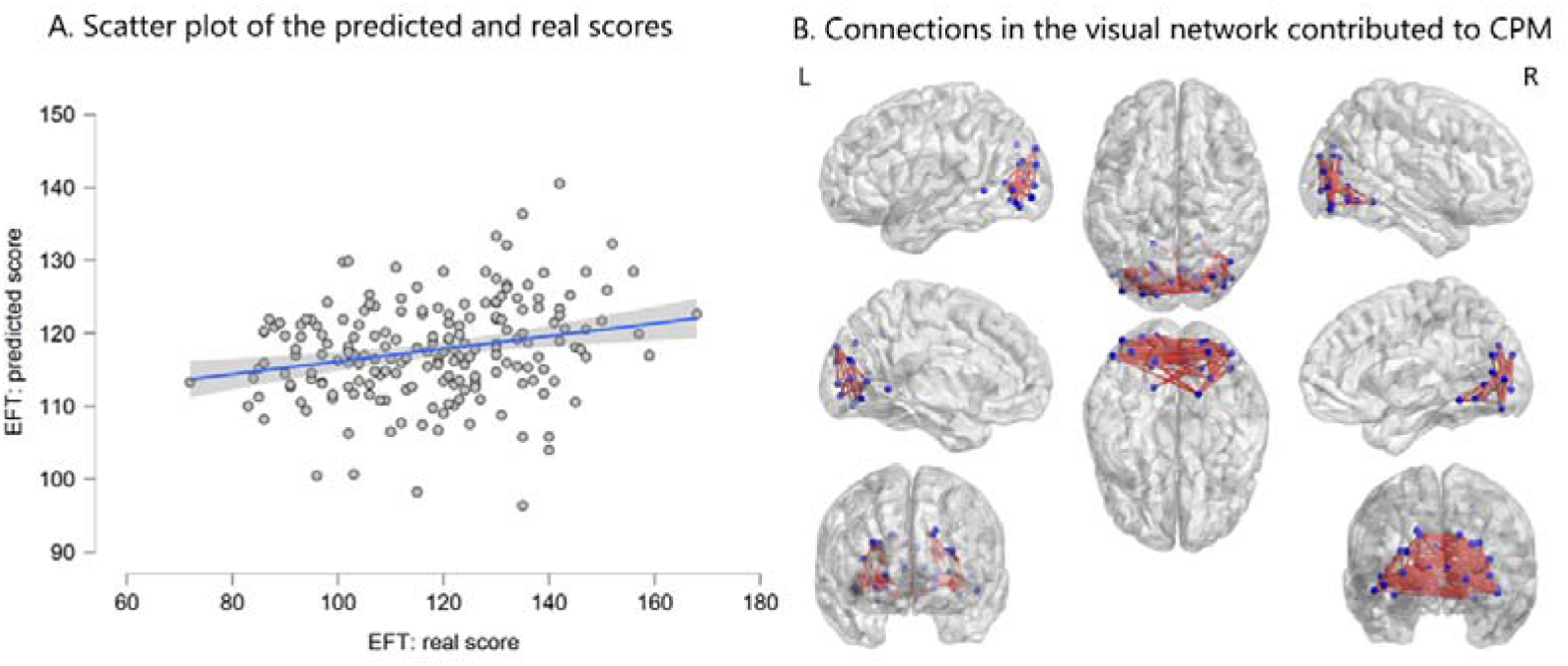
**(A)** Predicted score of EFT (CPM model; visual network parceled by Power-264 atlas; using 0.05 as feature selection threshold) was positively correlated with real score of EFT (*r*(191) = 0.24, p < 0.001), indicating good prediction performance of CPM model. Each dot represents one subject, and the gray area indicates 95% confidence interval for the linear fitting line. **(B)** Figure visualizing contribution of the connection in the visual network to CPM (parceled by Power-264 atlas; using 0.05 as feature selection threshold), thicker edge indicating more important to CPM.

**Table 1.** Prediction performance of CPM model on visual network using different parcellation schemas and feature selection threshold

| Feature selection threshold | Parcellation schema |  |  |
| --- | --- | --- | --- |
|  | Power-264 | Schaefer-400 | Zalesky-1024 |
| 0.1 | 0.24*** | 0.18* | 0.19** |
| 0.05 | 0.25*** | 0.18* | 0.20** |
| 0.01 | 0.24*** | 0.20** | 0.19** |

Given the important role of DMN on EFT, we additionally performed CPM analysis on DMN to predict EFT ability. We found that the prediction model of EFT generated from DMN was not significant (using 0.1, 0.05 and 0.01 as feature selection thresholds, p > 0.05), suggesting that intrinsic functional connectivity strength in DMN may not be the neural substrates underlying EFT ability, especially the sensory-perceptual qualities of EFT.

Table 1 The numbers represent the correlation coefficient between predicted score and real score, indicating prediction performance; \**p*< .05; \*\**p*< .01; \*\*\**p* < .001.

### 3.4 Global Efficiency of Visual Network Predicts EFT

To further explore the relationship between visual network and EFT, we adopted graph-analytical analysis and used topological property (global efficiency) of the visual network to predict EFT. Global efficiency of the visual network (using 0.2 as feature selection threshold) ranged from 0.2433 to 0.4382. Correlation analysis showed that global efficiency of the visual network is positively correlated with EFT-SPQ score (*r*(191) = 0.25, *p* < 0.001, using 0.2 as feature selection threshold). A linear model constructed with global efficiency predicted EFT-SPQ significantly (*r*(191) = 0.22, *p*< 0.001, using 0.2 as feature selection threshold and leave-one-subject-out cross-validation method, see Figure 3). The result was consistent across all thresholds (for correlation analysis, when using 0.1 or 0.3 as thresholds, *r*(191) = 0.26, *p* < 0.001; for predictive model performance, when using 0.1 or 0.3 as thresholds, *r*(191) = 0.22, *p* < 0.001). Collectively these results also robustly suggested that topological efficiency of visual network can predict EFT-SPQ, supporting it being the neural substrate of EFT ability.

**FIGURE 3.**
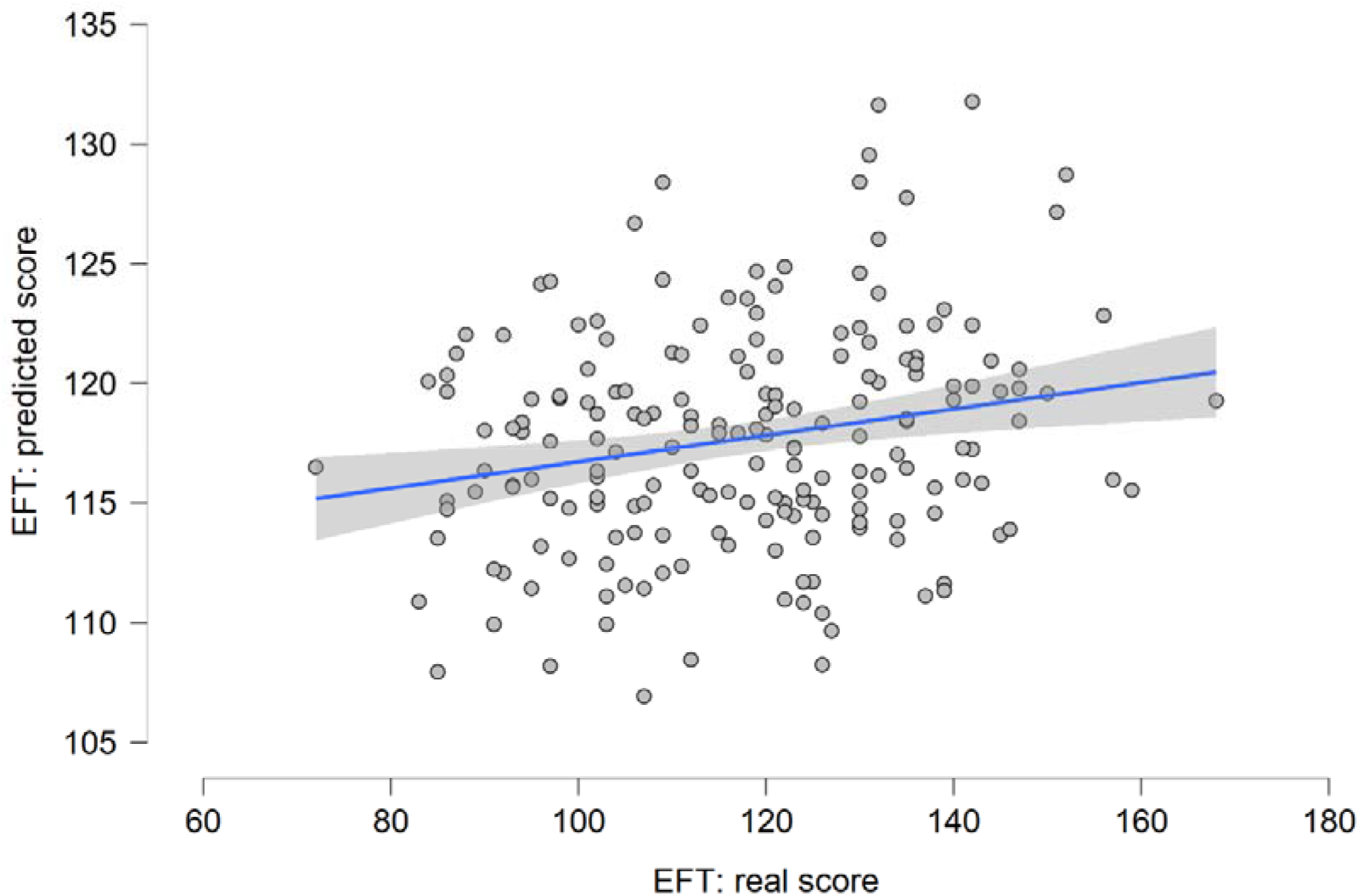
Predicted score of EFT (using network efficiency as predictor, 0.2 as feature selection threshold) was positively correlated with real score of EFT(*r*(191) = 0.24, p < 0.001), indicating good prediction performance of CPM model. Each dot represents one subject, and the gray area indicates 95% confidence interval for the linear fitting line.

Similar to CPM analysis, we analyzed the relationship between EFT and network efficiency of DMN. The correlation between global efficiency of DMN subnetworks and EFT-SPQ was not significant (using 0.2 as feature selection threshold; *r*(191) = 0.05, *p*> 0.05), thus we did not perform prediction model from global efficiency of DMN.

## 4 Discussion

In this study, we explored the neural substrates underlying the EFT ability through intrinsic functional connectivity, using CPM analysis and graph-theoretical analysis. Results from CPM analysis indicated that the imagining quality of sensory-perception in EFT can be predicted by the intrinsic functional connectivity strength in the visual network. Additionally, results from graph-theoretical analysis revealed that intrinsic topologic efficiency of the visual network is also a significant predictor of EFT ability. Collectively, these results supporting our hypothesis that visual network being the neural substrate underlying EFT ability, from a comprehensive perspectives of resting-state functional connectivity strength and the neural network.

This study revealed the relationship between the functional connectivity strength of visual network and the EFT ability, which is consistent with our hypothesis that visual network may serve as the neural basis underlying EFT ability. In addition, our findings converge with previous studies investigating the role of visual network in EFT, which have suggested that regions of the visual network such as the occipital cortex is involved during EFT (Bellana et al., 2016; Madore et al., 2016) and mental imagery (Dijkstra et al., 2017; Naselaris et al., 2015; Quoidbach et al., 2008). The finding is also supported by the behavioral studies about EFT, which indicated that guided eye movement would impair the quality of EFT (de Vito et al., 2015). Just as Conti and Irish (2021) pointed out that people subconsciously consider EFT as a form of visual activity, by using expressions such as “look ahead” or ‘‘see the future”. This inherently visual nature of EFT is supported by our finding. Theorists in the EFT area also suggested the visual nature of EFT at a neural level, postulated that memory and sensory-perceptual representations were constructed in the visual network, creating vivid imagery about future events (Conti & Irish, 2021; Pearson, 2019). Identical with these theories, present study discovered that the vividness of EFT can be predicted by the connectome strength of the visual network based on resting-state fMRI, thus the visual network could be the stable biomarker of EFT ability, indicating that people with well-functional visual network may perform EFT with more vivid details.

Apart from connectivity strength, this study further discovered that intrinsic network topological efficiency (network efficiency) of visual network was also a significant predictor of EFT ability, which indicated network organization of the visual network was equally important as connectivity strength, and might play an important role when forming vivid imagery about future events. With brain being a complex system, its function is considered the result of network organization (Bassett & Gazzaniga, 2011; Bullmore & Sporns, 2012b; Sporns & Betzel, 2016) rather than a simple summation of brain regions, thus the present study overcame such potential defect in CPM method by integrating it with graph-theoretical analysis. Global efficiency represents the efficiency of information transfer in the whole network, and higher global efficiency in brain network indicates higher information integration and processing ability of the brain (Latora & Marchiori, 2001), thus we considered that more efficient the visual network is, more vivid visual imagery about future events it will create owing to its finer function. On the other hand, this result also suggest that studies and theorists in EFT or mental imagery areas should not just focus on the cooperation’s of different brain functional subnetworks (Bellana et al., 2016; Conti & Irish, 2021; Pearson, 2019), but also the cooperation of subdivided components within a certain subnetwork, which may result in interesting findings about visual network’s activity during EFT. Overall, relationship between efficiency of visual network and EFT ability further validated the visual nature of EFT and expand it through the network perspective, suggesting that efficiency of information transfer in visual network also contributed to the variation of EFT ability.

The current finding did not find an association between functional connectome strength or network efficiency of DMN and EFT ability. Perhaps that is because our study focused on the vividness or the imagining quality of EFT, but not the other dimension of EFT about subjectively experiencing the constructed scenes, which may involve more in DMN since DMN is crucial for subjective consciousness (Crone et al., 2011; Fernández-Espejo et al., 2012). In future studies, the relationship between DMN and EFT ability should be investigated more comprehensively, especially the dimension about subjective experience (Autonoetic Consciousness), which may be affected by DMN function significantly.

In summary, the CPM method and graph-theoretical analysis was employed to reveal the relationship between visual network and EFT ability from a comprehensive perspective of resting-state functional connectivity strength and the neural network. The results indicated that the prediction models of EFT ability generated from connectome strength or network efficiency were both significant, jointly supporting visual network being the neural substrates underlying EFT ability and the hypothetical visual nature of EFT. The current finding provides preliminary evidence about the neural basis of EFT ability, especially its visual aspects. Given to the importance of visual function in EFT ability, it is worth exploring that whether training in visual imagery would improve one’s EFT ability (Abraham et al., 2018).

